# Survival and reinfection rates of SCTLD-affected corals treated in situ with amoxicillin

**DOI:** 10.1101/2025.06.18.660395

**Authors:** Karen L. Neely, Robert J. Nowicki, Michelle A. Dobler, Kathryn A. Toth, Kevin A. Macaulay, Sydney M. Gallagher

## Abstract

The unprecedented mortality to Caribbean corals caused by stony coral tissue loss disease (SCTLD) led to the use of an in-water medicine applied directly to disease lesions. This topical amoxicillin paste is highly effective in halting lesions and has been used on tens of thousands of wild corals since 2019, but long-term survival rates of treated corals as well as the frequency of potential reinfections remained speculative. We fate-tracked thousands of corals treated for SCTLD in the Florida Keys across numerous species and two habitats (inshore patch reefs and offshore spur and groove) every two months, assessing health condition and providing additional treatments if necessary. After three years, 84% of corals remained alive. Inshore corals had higher survival rates than offshore corals, and there were species-specific differences in survival, with the boulder corals *Montastraea cavernosa* and *Orbicella faveolata* having higher survivorship than brain coral species. Across all treated corals, 36% remained disease-free for at least one year after the initial treatment, and an additional 18% remained disease-free if any new lesions were treated within three months after the initial treatment. Reinfection rates were influenced by both habitat and species, with inshore corals more likely to remain disease-free than offshore corals. Among the species assessed, *Montastraea cavernosa* was the most likely to remain disease free, while the brain corals *Diploria labyrinthiformis* and *Colpophyllia natans* were the most prone to reinfection. These measurements can help guide expectations for disease intervention projects, including survival estimates if corals are regularly visited, as well as predictions of survivorship if diseased corals are visited only once or twice.

## Introduction

Since 2014, stony coral tissue loss disease (SCTLD) has had catastrophic impacts on scleractinian coral species in the Caribbean. First identified on corals off the coast of Miami, Florida (Precht et al. 2016), the disease spread both northward and southward along Florida’s Coral Reef, reaching the furthest stretches by the end of 2021. Beginning in 2017, SCTLD was also found outside of Florida, and has since spread to much of the Caribbean, affecting 33 countries/territories by June 2025 (Kramer et al. 2025). SCTLD is known to be contagious via physical contact and through the water column (Aeby et al. 2019, Meiling et al. 2021a). While spread within reef systems has generally followed oceanographic currents (Muller et al. 2020, Brandt et al. 2021, Estrada-Saldívar et al. 2021, Dobbelaere et al. 2022), its spread to different regions often has not, with shipping proposed as a possible vector for transmission across large distances (Dahlgren et al. 2021, Evans et al. 2022, Studivan et al. 2022).

SCTLD has exceptionally high infection and mortality rates on over 20 susceptible coral species (Precht et al. 2016, Hawthorn et al. 2024), including most of the primary reef-builders and 5 of the 7 Caribbean coral species listed under the US Endangered Species Act. Losses associated with SCTLD to date include:

1. 59% loss of coral cover in the Southeast Florida region (Hayes et al. 2022)
2. 46% loss of coral cover in Cozumel (Estrada-Saldívar et al. 2021)
3. 62% loss of coral cover in the Turks and Caicos (Heres et al. 2021)
4. Functional extinction of the pillar coral *Dendrogyra cylindrus* in Florida (Neely et al. 2021a)

Loss of corals has also led to significant declines in species richness, species diversity, reef functionality, and calcification rates (Estrada-Saldívar et al. 2020, Heres et al. 2021, Alvarez-Filip et al. 2022) on affected reefs.

SCTLD presents as focal or multi-focal lesions which progress across live coral tissue. Lesions often start at colony edges, but can also initiate in the middle of live coral tissue. Lesions are characterized by a generally sharp line of live tissue abutting bright white coral skeleton, though sometimes a layer of tissue on the edge can be seen separating from the skeleton. Histology of disease lesions has shown that necrosis originates at the basal body wall and appears to be associated with zooxanthellae (Landsberg et al. 2020). Despite much effort in assessing both the bacterial and viral communities of SCTLD-affected and unaffected colonies, no causative pathogen has yet been identified (Rosales et al. 2020, Clark et al. 2021, Work et al. 2021, Veglia et al. 2022, Rosales et al. 2023).

In response to the catastrophic losses caused by SCTLD, efforts were made to develop in-water coral medicines to treat and prevent mortality. Numerous laboratory and field efforts showed that natural products tested to date, UV light, and chlorinated epoxy are not effective for this purpose (Enochs and Kolodziej 2018, Neely 2018, Neely 2020, Neely et al. 2021b, Shilling et al. 2021). However, success was found through the development of a topical paste mixed with amoxicillin trihydrate (Neely et al. 2020, Neely et al. 2021b, Shilling et al. 2021, Walker et al. 2021a). The paste, termed Base2b (Ocean Alchemists), facilitates a three-day time release of the amoxicillin into the coral tissue. It is applied directly to active disease lesions by hand. The amoxicillin paste is largely effective at halting SCTLD lesions, with efficacy rates of 59 to 97% (Table 1).

**Table 1.** Summary of effectiveness of the amoxicillin and Base2b paste in halting active SCTLD lesions. Percentages of lesions halted and sample sizes (in parentheses) are shown, separated by species and study. For *Montastraea cavernosa*, some studies applied treatment only to the active lesion (no trench) while others added a treatment-filled firebreak (trench) a few centimeters from each active lesion.

| Source |  | Neely et al, 2020 | Neely et al, 2021b | Walker et al, 2021a | Walker et al, 2021b | Shilling et al, 2021 | Williams & Muller, 2025 |
| --- | --- | --- | --- | --- | --- | --- | --- |
| Location |  | Florida Keys | Florida Keys | SE Florida | SE Florida | SE Florida | St. John, USVI |
| *Montastraea cavernosa* | No trench | 89% (9) | 93% (287) | 59% (34) | 82% (11) |  | 60% (5) |
|  | Trench |  |  | 91% (34) |  | 95% (22) |  |
| *Orbicella faveolata* |  | 91% (23) | 93% (848) |  | 82% (177) |  |  |
| *O. franksii* |  |  |  |  |  |  | 100% (5) |
| *O. annularis* |  |  | 93% (15) |  |  |  | 100% (15) |
| *Colpophyllia natans* |  | 67% (3) | 94% (98) |  |  |  |  |
| *Diploria labyrinthiformis* |  | 88% (8) | 75% (51) |  |  |  |  |
| *Pseudodiploria strigosa* |  | 73% (15) | 97% (399) |  |  |  | 70% (10) |
| *P. clivosa* |  |  | 94% (37) |  |  |  |  |
| *Dichocoenia stokesii* |  |  | 95% (102) |  |  |  |  |
| *Solenastrea bournoni* |  |  | 100% (12) |  |  |  |  |
| *Siderastrea siderea* |  |  | 90% (188) |  | 100% (9) |  |  |

Despite the success of Base2b in halting existing legions, treatments are not prophylactic, and new lesions can develop on previously treated colonies. To date, the rates and patterns of reinfection and survival of previously treated colonies are not well quantified. This inhibits practitioners from estimating the long-term efficacy of past interventions, as well as optimizing intervention operations. To characterize these patterns, we treated and monitored over 5000 coral colonies representing 19 coral species in the Florida Keys National Marine Sanctuary on an approximately eight week cadence from January 2019 to July 2024. These efforts allowed us to fate-track treated colonies to assess reinfection and survival rates.

## Methods

### Treatment Sites

We assessed amoxicillin-treated corals from three offshore sites and four inshore sites on Florida’s Coral Reef (Table 2; Figure 1). The offshore sites were 7 – 10.5 km from land, had spur-and-groove morphology, and had corals treated at depths of 2.0 – 9.5 meters. In contrast, the inshore sites were 0.6 – 2.9 km from land, were patch reefs surrounded by sand and seagrass, and had corals treated at depths of 1.8 – 6.0 meters. Inshore reefs had higher coral density and coral species diversity than offshore reefs. Reefs were first visited and treated between January 2019 and August 2021. Time of treatment was based on when permits were authorized as well as when SCTLD outbreaks were observed. Once a reef was initially treated, it was revisited approximately one month afterwards, and then approximately every two months after that for full surveys and treatments if necessary.

**Table 2.**
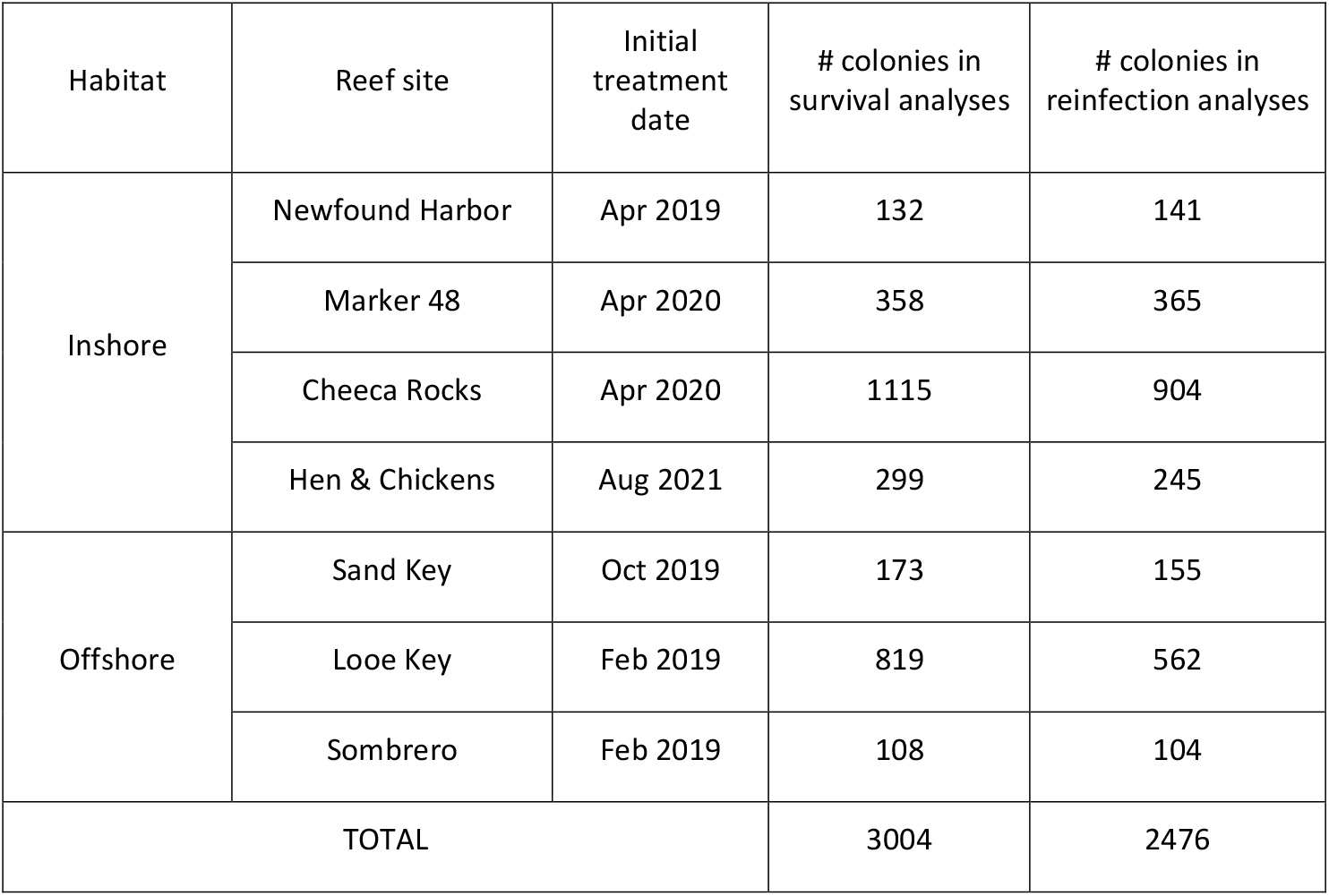
Initial treatment date and number of colonies used in the two fate-tracking analyses (survivorship curves and reinfection patterns) across each treatment site.

| Habitat | Reef site | Initial treatment date | # colonies in survival analyses | # colonies in reinfection analyses |
| --- | --- | --- | --- | --- |
| Inshore | Newfound Harbor | Apr 2019 | 132 | 141 |
|  | Marker 48 | Apr 2020 | 358 | 365 |
|  | Cheeca Rocks | Apr 2020 | 1115 | 904 |
|  | Hen & Chickens | Aug 2021 | 299 | 245 |
| Offshore | Sand Key | Oct 2019 | 173 | 155 |
|  | Looe Key | Feb 2019 | 819 | 562 |
|  | Sombrero | Feb 2019 | 108 | 104 |
| TOTAL |  |  | 3004 | 2476 |

**Figure 1.**
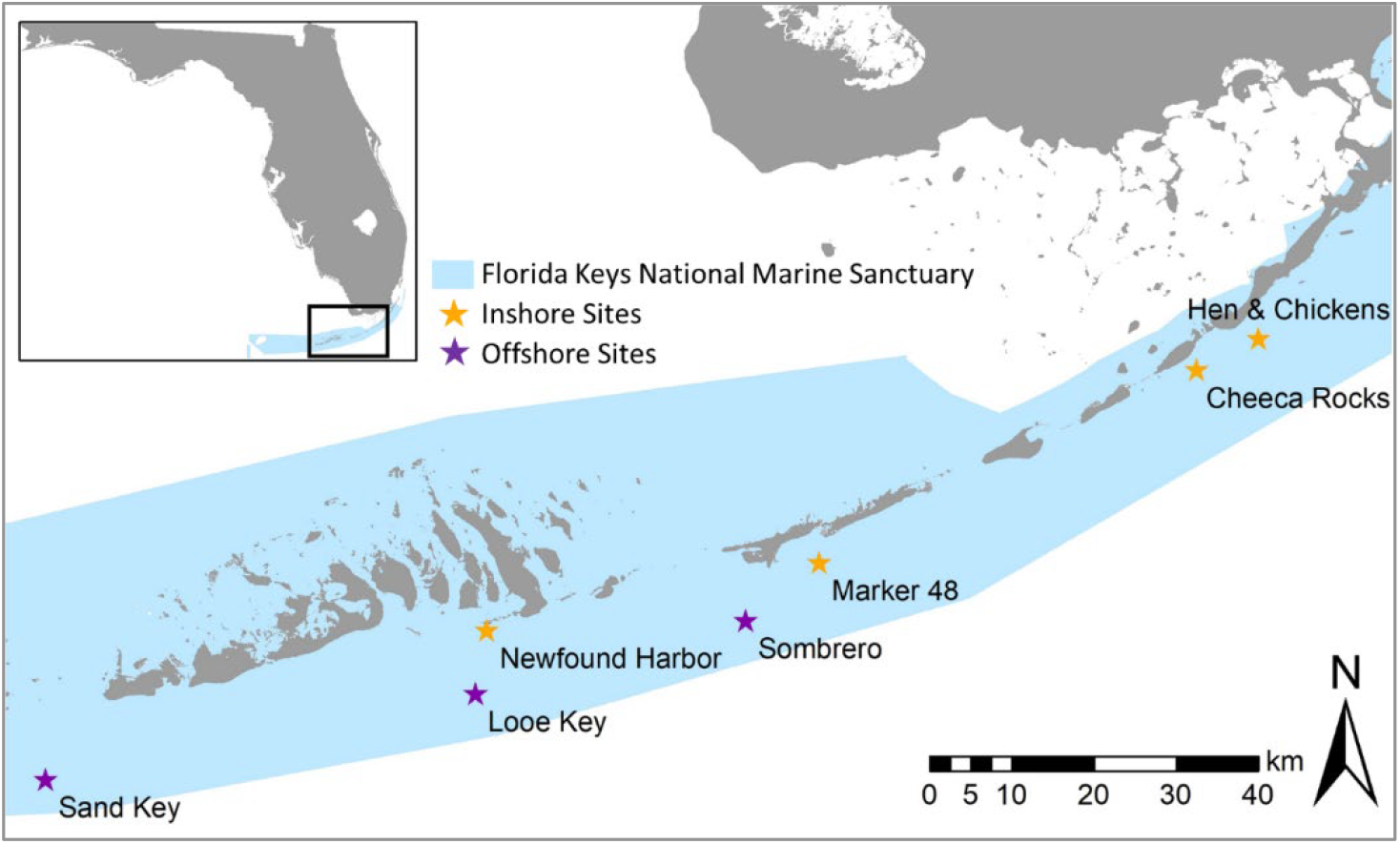
Map of assessed intervention sites within the Florida Keys National Marine Sanctuary. Sites included three offshore forereef locations (purple stars) and four inshore patch reef locations (orange stars).

### Treatments and Monitoring

When an SCTLD-affected coral was found, a numbered cattle tag was affixed for future identification. The coral was also mapped and photographed from multiple angles both before and after treatments were applied. All active SCTLD lesions were treated with 98% amoxicillin trihydrate powder hand-mixed in an 8:1 Base2b:amoxicillin by weight ratio. Mixing was done manually between 1 and 72 hours before application to the diseased corals. The mixed paste was packed into 60cc catheter syringes, then were squeezed out in a ∼1 cm wide band along the active lesion. The paste was pressed by hand onto the colony so that approximately 0.5 cm of treatment covered the live but diseased tissue and about 0.5 cm was pressed into the recently denuded skeleton to anchor the treatment onto the colony. In rare occasions where the treatment did not adhere well, such as in surge or occasionally on excessively mucoused corals, small (∼ 0.25 cm) pieces of modeling clay were placed sporadically atop the treatment to anchor it to the coral.

Divers used maps and datasheets with coral tag numbers, species, and directions to find each previously-tagged coral. From February 2019 to February 2022, every coral was visited approximately every two months for a health assessment and retreatment if needed. Each coral was coded as either “T” (treated), “NAD” (no active disease), “DEAD” (newly dead since the last monitoring event), or, rarely, “AL” (active lesions – not treated; this occurred only when the amount of remaining tissue was so small as to not be worth treatment efforts). Beginning in March 2022, methods shifted to allow for broader SCTLD detection and treatment coverage. Two sites – Sombrero Reef (offshore) and Cheeca Rocks (inshore) – retained the established level of intense colony-specific monitoring. At all other sites, divers used a systematic swim pattern to cover the entire reef to detect new lesions while still locating and assessing all previously tagged corals, but live corals without active SCTLD did not receive full monitoring. Corals which did have SCTLD were recorded as T or AL, and any dead corals were recorded as such. At all sites, any newly diseased corals found during each monitoring period were tagged, treated, and added to all future monitoring events. Monitoring time steps were rounded to the nearest number of 30 day months since initial treatment (e.g. a visit 14-45 days after initial treatment was rounded to 1 month, 46-75 rounded to 2 months, etc.).

### Survival rates

To assess overall coral survivorship rates, we used the *survfit* function in the *survival* package (Therneau 2024) to fit a global Kaplan-Meyer survival curve using the number of days observed and survivorship. The monitoring data were uninformative right-censored (i.e. corals were followed until they either died or the study ended). We focused on monitoring data for five species for which suitable replicates were available in both habitats: *Orbicella faveolata* (n = 1079), *Montastraea cavernosa* (n = 708), *Pseudodiploria strigosa* (n = 163), *Colpophyllia natans* (n = 802), and *Diploria labyrinthiformis* (n = 252). The date each coral was first observed dead was considered its date of death, which may have slightly overestimated survival rates for some corals by up to two months. Assessments of mortality for most sites were conducted from the initial treatment date (Table 2) through June 30, 2023. For Newfound Harbor, assessments past May 30, 2022 were excluded because a presumably different disease swept through *Pseudodiploria clivosa* colonies at the site, thus confounding SCTLD survival data. We also generated survival curves for each species across each habitat. To assess the effects of coral species and habitat (inshore, offshore) on coral survival, we performed a series of log-rank tests using the *survdiff* function in the *survival* package (Therneau and Grambsch 2000, Therneau 2022) in R version 4.2.2 (R Core Team 2022) followed by pairwise log-rank comparisons using the *pairwise_survdiff* function in the *survminer* package. The false discovery rate (FDR; Benjamini and Hochberg (1995)) was used to correct for multiple comparisons.

### Reinfection patterns

Health status and reinfection rates through time were assessed for all corals that met the following conditions:

1. Were from one of the five species most commonly treated across inshore and offshore reefs: *Orbicella faveolata, Montastraea cavernosa, Pseudodiploria strigosa, Colpophyllia natans*, and *Diploria labyrinthiformis*.
2. Had been visited at least once during each quarter after the initial treatment (1-3 months, 4-6 months, 7-9 months, and 10-12 months). As such, only corals first treated before July 2023 were included.
3. Did not have any partial or full-mortality related to the 2023 marine heatwave (Neely et al. 2024).

For corals with more than one data point per quarter, we used the most detrimental condition as the point of record for that time point: Dead > Treated > Active lesion untreated > No active disease. We then grouped corals by each health status through each time point, and calculated the proportion that were lesion-free from each time point through the end of the first year since initial treatment.

A total of 2,476 corals were assessed for 1-year reinfection patterns after the initial treatment date (Table 2). These were from the five most common species at both inshore and offshore sites, with 1,655 from inshore sites and 821 from offshore sites. To assess how 1) species and 2) habitat (inshore / offshore) affected the likelihood of a coral remaining disease-free for at least one year after one visit (initial treatment only) or two visits (initial treatment + an additional visit and treatment if needed at 1-3 months), we created four generalized linear models (GLMs): one with species only, one with habitat only, one with both but without an interaction effect, and a final ‘full’ model that also included an interaction between species and habitat (Supplemental Table 1). We used a model selection approach (Anderson 2007) leveraging Akaike’s Information Criterion (AIC; Akaike (1973)), to choose the optimal model for inference, selecting the model with the lowest AIC score (and thus best fit). This model was used to assess whether habitat or species were significant predictors of reinfection rates. We assessed predictive power by splitting the data into a 70/30 train/test split measuring area under the curve (AUC) of the predictive values. Statistical analyses were conducted in R v.4.4.1 (R Core Team, 2022) using the *stats* package for generalized linear models as well as the packages *ROCR* for the AUC (Sing et al. 2005), *emmeans* for post hoc analysis (Lenth 2018) and *multcomp* for group significance designations (Hothorn et al. 2008).

## Results

### Survival rates

Overall coral survivorship as assessed by the global Kaplan-Meyer curve was 92% at one year, 89% at two years, and 86% at three years. Survival rates did, however, vary between habitats (inshore vs. offshore) and across species (Figure 2).

**Figure 2.**
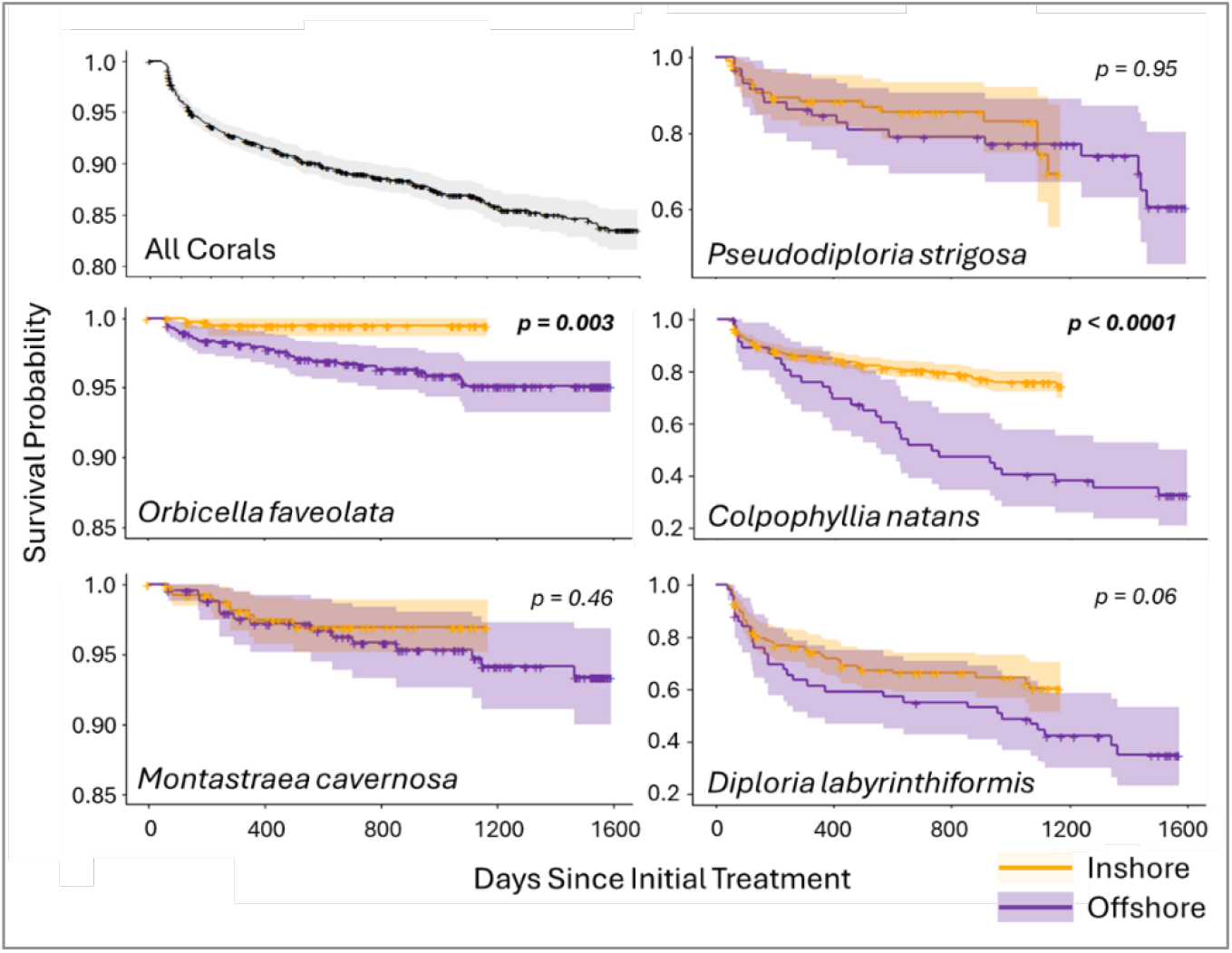
Kaplan-Meier survival curves for corals regularly monitored and treated if needed. Survival probabilities are shown for all five coral species combined as well as each species across both inshore (orange) and offshore (purple) habitats. Shaded areas represent confidence intervals, and crosses indicate censored corals (those for which monitoring terminated and the individual was still alive). Species for which inshore and offshore habitat survival rates were significantly different have p values shown in bold.

Survival differed significantly by both habitat (χ^2^ (3004) = 15.94, p = 0.0001) and species (χ^2^ (3004) = 447.69, p < 0.0001) when considered globally. The impact of habitat on survivorship was mediated by species. Specifically, differences in survival rates between habitats were driven by two species – *O. faveolata* (χ^2^ (1079) = 8.93, p = 0.003) and *C. natans* (χ^2^ (804) = 26.59, p < 0.0001) – which both had higher survivorship inshore than offshore (Figure 2). The other three species did not have significantly different survivorship between habitat types, though *D. labyrinthiformis* trended towards significance (χ^2^ (252) = 3.44, p = 0.06). Survivorship of all species at inshore reef sites remained above 60% through 1200 days (3.25 years). At offshore reef sites, *O. faveolata, M. cavernosa*, and *P. strigosa* had survival rates above 60% at 1600 days (4.4 years), but only about 1/3 of *C. natans* and *D. labyrinthiformis* survived that long.

Survivorship between species differed significantly, both globally and within habitat types (Figure 3). Specifically, at inshore sites, *O. faveolata* and *M. cavernosa* had significantly higher survivorship than all other species (all p < 0.00001; see Supplemental Table 2 for full statistical values), while *D. labyrinthiformis* had significantly lower survivorship than all other species (all p < .005). *Pseudodiploria strigosa* and *C. natans* had significantly lower survivorship than the top two performing species, and significantly higher survivorship than *D. labyrinthiformis*, but did not differ from each other. At offshore sites, *O. faveolata* and *M. cavernosa* had significantly higher survivorship than all other species (all p < 0.0001), and *C. natans* and *D. labyrinthiformis* had significantly lower survivorship than all other species (all p < 0.002). *Pseudodiploria strigosa* had significantly lower survivorship than the top two performing species and significantly higher survivorship than the bottom two. (Figure 3).

**Figure 3.**
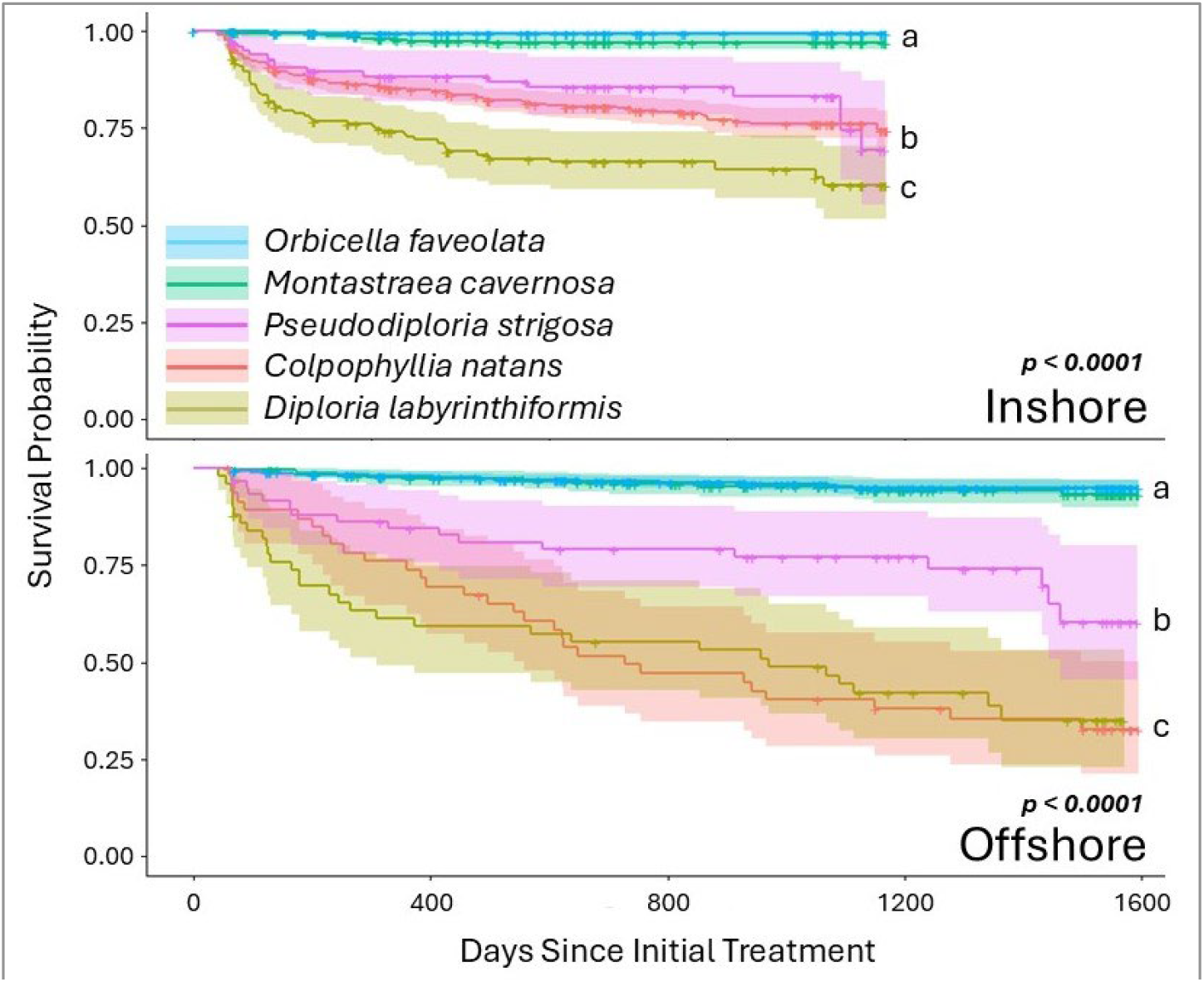
Kaplan-Meier survival curves for each of the two habitats, broken out by species. Shaded areas represent confidence intervals, and crosses indicate censored corals. For both habitats, *Orbicella faveolata* and *Montastraea cavernosa* had higher survivorship than the three brain coral species.

### Reinfection Patterns

Across all individuals, 36% (897/2476) remained disease free for at least one year after the initial treatment. An additional 18% (for a total of 54% (1336/2476)) remained disease free through at least the first year when new lesions, if present, were treated during the 1-3 month follow-up. The remaining 46% of corals all developed additional lesions after the initial 1-3 months, although the percentage of corals remaining disease-free continued to rise through the first year (Figure 4).

**Figure 4.**
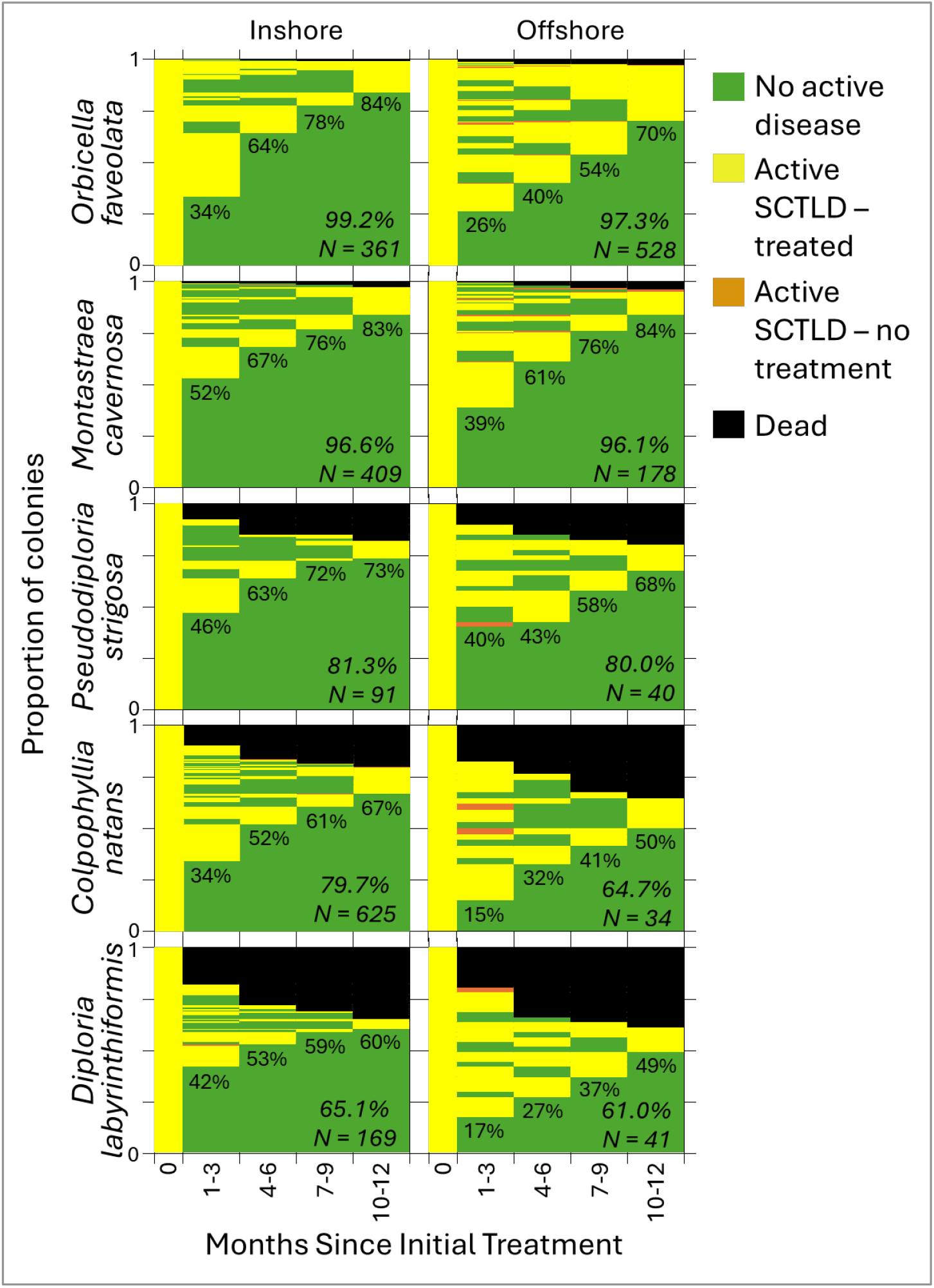
The proportion of coral colonies (separated by habitat and species) exhibiting each health condition during each monitoring period. The percentage of corals that were healthy and remained healthy through the first year is shown for each time point. The survival rate and sample size for each species/habitat combination is shown in italics.

For comparing habitat and species effects on corals remaining disease-free after only the initial treatment, the best model included both variables but no interaction factor (Supplemental Table 1). Both habitat (p < 0.001) and species (p < 0.001) were significant variables, though the model was not particularly predictive (AUC = 0.61).

At inshore reef sites, 36% of assessed corals remained disease free after the initial treatment, compared to only 29% of offshore corals (Figure 5). Globally (across inshore and offshore habitats combined), 48% of *M. cavernosa* remained disease free, significantly more than *C. natans* (33%; p < 0.0001), *O. faveolata* (29%; p < 0.0001), and *D. labyrinthiformis* (37%; p = 0.02). Additionally, significantly more *P. strigosa* remained disease free (44%) than *C. natans* (33%; p = 0.02). Within both inshore and offshore habitats, *M. cavernosa* colonies were less likely to develop lesions than both *C. natans* and *O. faveolata*. When comparing species and habitats together, every one of the five species was less likely to develop additional lesions within the inshore habitat compared to the offshore after initial treatment (all p < 0.0001). Additionally, offshore *O. faveolata* were more likely to develop lesions than every inshore species (all p < 0.02; Supplemental Table 3), offshore *C. natans* were more likely to develop lesions than every inshore species except *O. faveolata* (all p < 0.01), and offshore *D. labyrinthiformis* were more likely to develop lesions than inshore *P. strigosa* and *M. cavernosa* (both p < 0.03).

**Figure 5.**
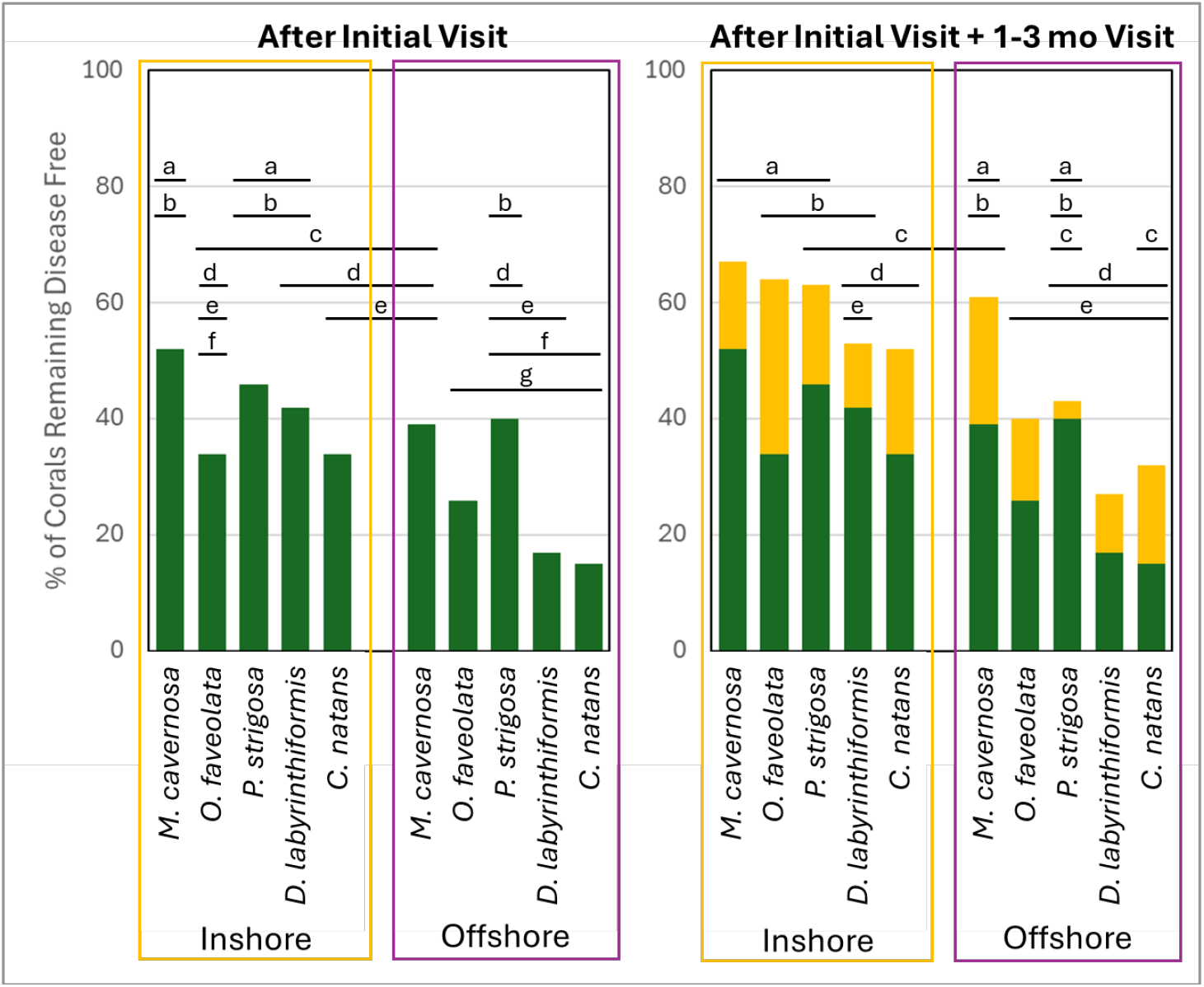
The proportion of coral colonies (separated by habitat and by species) that remained disease-free for at least one year after the initial treatment (green only) or after the initial treatment plus an additional treatment at 1-3 months if needed (green + yellow). Letters above the bars indicate percentages that are not significantly different from each other

For assessing the likelihood of a coral remaining disease free for at least one year after the initial treatment plus a retreatment if necessary at 1-3 months, the best model included habitat, species, and the interaction of factors. Again, this model was not particularly predictive (AUC = 0.61). However, habitat was a significant variable, with 55% of inshore reefs remaining disease free compared to 44% of offshore reefs (p < 0.001; Figure 5). Species was also a significant predictor (p < 0.001), as was the interaction between species and habitat (p = 0.03). On inshore reefs, 67% of *M. cavernosa* remained disease-free after the initial plus 1-3 month checkups, a value significantly greater than for *C. natans* (52%; p < 0.0001) and *D. labyrinthiformis* (53%; p = 0.03). Also on inshore reefs, significantly more *O. faveolata* remained disease-free (64%) than *C. natans* (52%; p = 0.007). On offshore reefs, 40% of *O. faveolata* and 61% of *M. cavernosa* remained disease free, values significantly higher than for *D. labyrinthiformis* (27%; p < 0.001 and p = 0.006 respectively). When comparing species and habitats together, inshore *O. faveolata* and *M. cavernosa* were significantly more likely to remain disease-free than offshore *C. natans, D. labyrinthiformis*, and *O. faveolata* (all p < 0.02) Inshore *P. strigosa* were significantly more likely to remain disease-free than offshore *D. labyrinthiformis* and *O. faveolata* (both p < 0.01). And inshore *C. natans* were significantly more likely to remain disease-free than offshore *O. faveolata* (p = 0.001; Supplemental Table 3).

## Discussion

Survivorship varied significantly by species in both habitats, with obvious “winners” and “losers”. The boulder corals, *O. faveolata* and *M. cavernosa*, had significantly higher survival than the brain corals, *C. natans, D. labyrinthiformis*, and *P. strigosa*, in both habitats (Figure 3). In contrast, the importance of habitat on survivorship depended on species, particularly for *O. faveolata* and *C. natans* (Figure 2). At an assemblage level (when combining all 5 species), survivorship was higher for inshore than offshore reef sites. This was consistent regardless of whether the coral had high survival rates (e.g. *O. faveolata*) or low survival rates (e.g. *C. natans*).

Reinfection rates by species roughly match definitions of SCTLD susceptibility among species. *Montastraea cavernosa* was the species least likely to reinfect at both inshore and offshore habitats, while *D. labyrinthiformis* and *C. natans* were most likely to reinfect. The case definition for SCTLD likewise defines these two brain coral species as highly susceptible, and *M. cavernosa* as moderately susceptible (Hawthorn et al. 2024). Just as corals vary in their susceptibility to their initial infections, subsequent infections follow similar patterns. The high survival rates of treated *M. cavernosa* and *O. faveolata* are likely somewhat attributable to their lower susceptibility and reinfection rates. However, we also anecdotally noted other characteristics that improved a colony’s likelihood of survivorship under the two-month visitation regime: colony size, lesion progression rate, and continuity of tissue. Colonies that were small, had continuous tissue, and had rapid lesion progression rates (characteristics more common on the brain coral species; see Meiling et al. (2020) and Meiling et al. (2021b) for progression rates) were more likely to have an SCTLD lesion consume all available tissue in the time between visitations. In contrast, colonies that were large, had multiple tissue isolates, and had slower lesion progression rates (characteristics more common on the boulder coral species) were more likely to be visited and treated before full mortality occurred, or to have non-diseased tissue isolates remaining. Left untreated over time, we suggest that mortality of *M. cavernosa* and *O. faveolata* could approach that of the brain coral species.

It is not certain why inshore corals had lower infection rates and higher survivorship than offshore corals, but there are notable differences between the habitat types. One likely factor is the greater temperature extremes at inshore reef sites compared to offshore, including coral paling and/or bleaching at inshore reef sites every summer during the study. Corals at offshore reef sites did not pale or bleach during any summer of the survey time period. Coral bleaching is known to slow and/or halt SCTLD lesions (Meiling et al. 2020), and this annual event is likely to have assisted intervention efforts at inshore reef sites by halting new lesion developments and keeping pathogen load under control. As suggested in Neely et al. (2021b), the isolation of the inshore patch reefs from other coral areas via sand and seagrass meadows may have also lowered reinfection rates by slowing the reintroduction of pathogenic particles after a bleaching event, thus providing protection from transmission and new lesion development even months after a bleaching event. We suggest that the cessation of infection associated with bleaching is likely to be meaningful at landscape scales and ecological timelines.

Though amoxicillin treatments are known to be highly effective at halting lesions, they are not prophylactic. New lesions can and do develop on treated colonies. However, the extent of additional lesion development was previously unknown, and thus anticipating the level of commitment to treatments on corals to keep them alive was also not well defined. Our work here shows the value of consistent visitations in maintaining a high survival rate, and also identifies that single- or double-visitations can prevent mortality on a large proportion of treated corals. On offshore reefs in the Florida Keys, we show that, depending on species, between 15% and 40% of corals treated once will not develop additional lesions for at least one year, with *P. strigosa* and *M. cavernosa* performing the best. If corals were visited twice – for an initial visit and a follow-up at 1-2 months, between 27% and 61% of offshore corals (variation by species) would remain disease-free, while at inshore reefs, one- or two-time visitation success rates would vary be 52-72% (variation by species). Because the efficacy of treatment visits has species and sometimes habitat-specific variation, each unit of treatment effort is likely to yield a different return depending on the target species and location. But, we show that regular visitation over long periods of time can keep most individuals alive, and that even single or double visitations, as may be more feasible in logistically challenging locations or in resource-limited scenarios, are likely to preserve substantial portions of these susceptible species’ populations.

## Supplemental Tables

**Supplemental Table 1:**
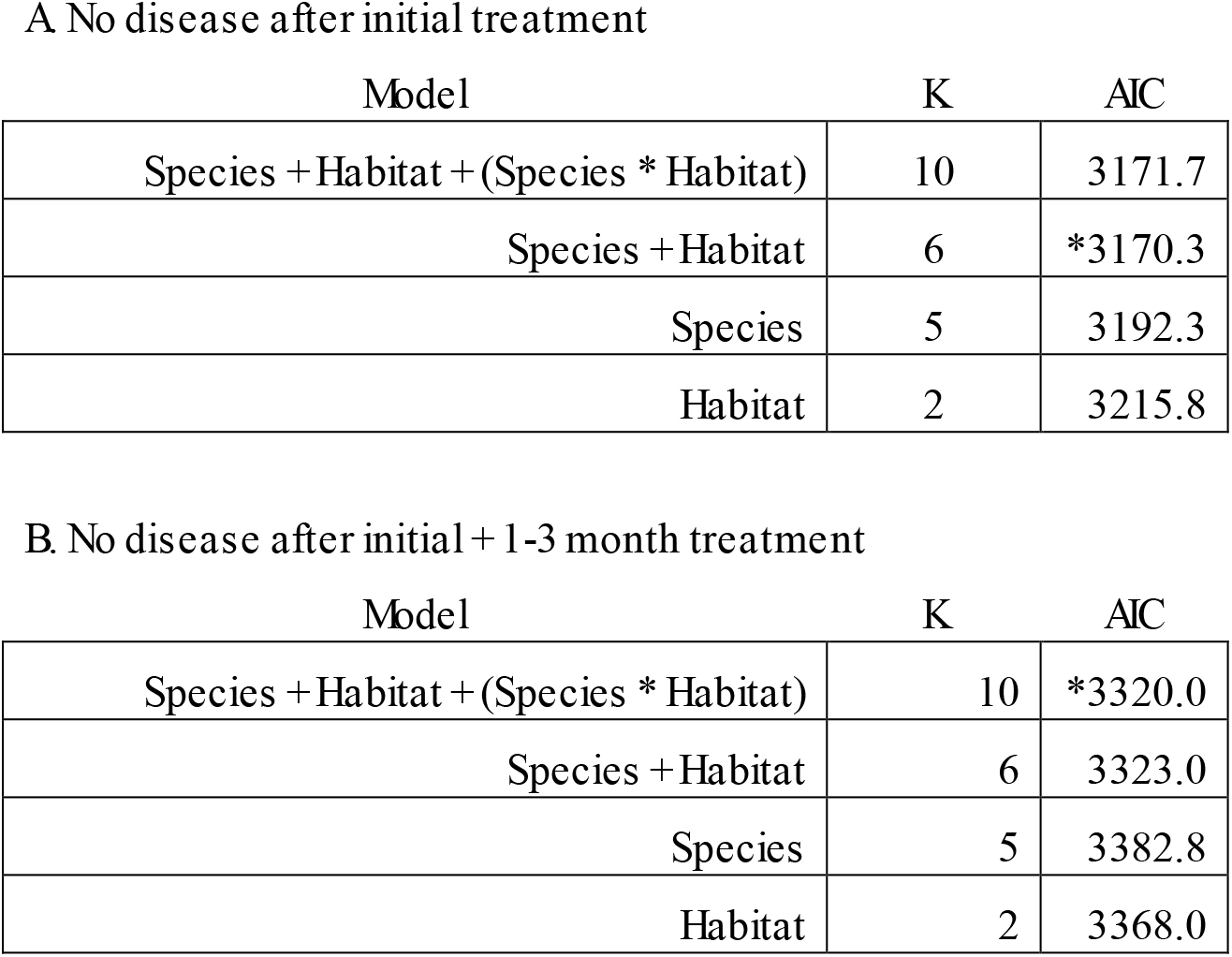
Number of parameters (K) and Akaike Information Criterion (AIC) for the models assessing the probability of SCTLD reinfection for at least one year after only an initial treatment (A), or after the initial treatment plus an additional visitation and treatment if needed at 1-3 months (B). Variables are species and habitat (inshore versus offshore). The models used are indicated with “*”.

A. No disease after initial treatment
| Model | K | AIC |
| --- | --- | --- |
| Species + Habitat + (Species * Habitat) | 10 | 3171.7 |
| Species + Habitat | 6 | *3170.3 |
| Species | 5 | 3192.3 |
| Habitat | 2 | 3215.8 |

B. No disease after initial + 1-3 month treatment
| Model | K | AIC |
| --- | --- | --- |
| Species + Habitat + (Species * Habitat) | 10 | *3320.0 |
| Species + Habitat | 6 | 3323.0 |
| Species | 5 | 3382.8 |
| Habitat | 2 | 3368.0 |

**Supplemental Table 2:** Statistical comparisons of Kaplan-Meier survival curves between species among inshore and offshore reefs. Significant differences are p-values shown in green. Species are identified as: OFAV (*Orbicella faveolata*), MCAV (*Montastraea cavernosa*), PSTR (*Pseudodiploria strigosa*), CNAT (*Colpophyllia natans*), and DLAB (*Diploria labyrinthiformis*).

|  | Inshore |  |  |  |  |
| --- | --- | --- | --- | --- | --- |
|  | OFAV | MCAV | PSTR | CNAT | DLAB |
| OFAV |  | 0.25 | <0.00001 | <0.00001 | <0.00001 |
| MCAV |  |  | <0.00001 | <0.00001 | <0.00001 |
| PSTR |  |  |  | 0.5 | 0.005 |
| CNAT |  |  |  |  | <0.0001 |
| DLAB |  |  |  |  |  |

Offshore
|  | Offshore |  |  |  |  |
| --- | --- | --- | --- | --- | --- |
|  | OFAV | MCAV | PSTR | CNAT | DLAB |
| OFAV |  | 0.55 | <0.00001 | <0.00001 | <0.00001 |
| MCAV |  |  | <0.00001 | <0.00001 | <0.00001 |
| PSTR |  |  |  | 0.002 | 0.002 |
| CNAT |  |  |  |  | 0.99 |
| DLAB |  |  |  |  |  |

**Supplemental Table 3:** The percentages of coral colonies that were healthy for at least one year following an initial treatment (left) and an initial treatment plus a 1-3 month monitoring and retreatment if necessary (right). Percentages follow the species codes, and significant differences are shown as green p-values. Tables compare (top to bottom) species within the inshore habitat, the offshore habitat, and between the inshore and offshore habitats. For the initial treatment comparisons, the interaction factor was not significant, and so the bottom panel shows species comparisons when both habitats are combined. Species are identified as: CNAT (*Colpophyllia natans*), DLAB (*Diploria labyrinthiformis*), OFAV (*Orbicella faveolata*), PSTR (*Pseudodiploria strigosa*), and MCAV (*Montastraea cavernosa*).

|  |  | Inshore |  |  |  |  |
| --- | --- | --- | --- | --- | --- | --- |
|  |  | CNAT (34%) | DLAB (42%) | OFAV (34%) | PSTR (46%) | MCAV (52%) |
| Inshore | CNAT (34%) |  |  |  |  | <.0001 |
|  | DLAB (42%) |  |  |  |  |  |
|  | OFAV (34%) |  |  |  |  | <.0001 |
|  | PSTR (46%) |  |  |  |  |  |
|  | MCAV (52%) |  |  |  |  |  |

|  |  | Offshore |  |  |  |  |
| --- | --- | --- | --- | --- | --- | --- |
|  |  | CNAT (15%) | DLAB (17%) | OFAV (26%) | PSTR (40%) | MCAV (39%) |
| Offshore | CNAT (15%) |  |  |  |  | <.0001 |
|  | DLAB (17%) |  |  |  |  |  |
|  | OFAV (26%) |  |  |  |  | <.0001 |
|  | PSTR (40%) |  |  |  |  |  |
|  | MCAV (39%) |  |  |  |  |  |

|  |  | Inshore |  |  |  |  |
| --- | --- | --- | --- | --- | --- | --- |
|  |  | CNAT (34%) | DLAB (42%) | OFAV (34%) | PSTR (46%) | MCAV (52%) |
| Offshore | CNAT (15%) | <.0001 | 0.01 |  | 0.0001 | <.0001 |
|  | DLAB (17%) |  | <.0001 |  | 0.03 | <.0001 |
|  | OFAV (26%) | 0.02 | 0.005 | <.0001 | <.0001 | <.0001 |
|  | PSTR (40%) |  |  |  | <.0001 |  |
|  | MCAV (39%) |  |  |  |  | <.0001 |

|  |  | All |  |  |  |  |
| --- | --- | --- | --- | --- | --- | --- |
|  |  | CNAT (33%) | DLAB (37%) | OFAV (29%) | PSTR (44%) | MCAV (48%) |
| All | CNAT (33%) |  |  |  | 0.02 | <.0001 |
|  | DLAB (37%) |  |  |  |  | 0.02 |
|  | OFAV (29%) |  |  |  |  | <.0001 |
|  | PSTR (44%) |  |  |  |  |  |
|  | MCAV (48%) |  |  |  |  |  |

%healthy following 1-3 month monitor/treatment
|  |  | Inshore |  |  |  |  |
| --- | --- | --- | --- | --- | --- | --- |
|  |  | CNAT (52%) | DLAB (53%) | OFAV (64%) | PSTR (63%) | MCAV (67%) |
| Inshore | CNAT (52%) |  |  | 0.007 |  | <.0001 |
|  | DLAB (53%) |  |  |  |  | 0.03 |
|  | OFAV (64%) |  |  |  |  |  |
|  | PSTR (63%) |  |  |  |  |  |
|  | MCAV (67%) |  |  |  |  |  |

|  |  | Offshore |  |  |  |  |
| --- | --- | --- | --- | --- | --- | --- |
|  |  | CNAT (32%) | DLAB (27%) | OFAV (40%) | PSTR (43%) | MCAV (61%) |
| Offshore | CNAT (32%) |  |  |  |  |  |
|  | DLAB (27%) |  |  | <.0001 |  | 0.006 |
|  | OFAV (40%) |  |  |  |  |  |
|  | PSTR (43%) |  |  |  |  |  |
|  | MCAV (61%) |  |  |  |  |  |

|  |  | Inshore |  |  |  |  |
| --- | --- | --- | --- | --- | --- | --- |
|  |  | CNAT (52%) | DLAB (53%) | OFAV (64%) | PSTR (63%) | MCAV (67%) |
| Offshore | CNAT (32%) |  |  | 0.02 |  | 0.005 |
|  | DLAB (27%) |  |  | 0.0007 | 0.009 | 0.0001 |
|  | OFAV (40%) | 0.001 |  | <.0001 | 0.003 | <.0001 |
|  | PSTR (43%) |  |  |  |  |  |
|  | MCAV (61%) |  |  |  |  |  |

## Author Contributions

KLN secured funding, designed the study, and managed the project. RJN and KAT conducted statistical analyses. KLN, MAD, KAC, and SMG collected data. MAD conducted data entry and data management. KAC managed photographic data. SMG managed GIS and field datasheets. KLN and RJN wrote the manuscript, and MAD, KAT, KAM, and SMG provided edits.

## Acknowledgments

In addition to the authors, a variety of lab members and collaborators assisted in coral treatments and monitoring; we are grateful to Force Blue, Emily Hower, Kayla Merrill, Nicole Charnock, Hailey Vaughan, and Arelys Chaparro. Work was conducted under Florida Keys National Marine Sanctuary permits 2018-141, 2019-115, and 2020-077.

## Data Availability Statement

The raw data supporting the conclusions of this article will be made available by the authors, without undue reservation.

## References

Aeby, G., B. Ushijima, J. E. Campbell, S. Jones, G. Williams, J. L. Meyer, C. Hase, and V. Paul. 2019. Pathogenesis of a tissue loss disease affecting multiple species of corals along the Florida Reef Tract. Frontiers in Marine Science 6.

Akaike, H. 1973. Maximum likelihood identification of Gaussian autoregressive moving average models. Biometrika 60:255–265.

Alvarez-Filip, L., F.J. González-Barrios, E. Pérez-Cervantes, A. Molina-Hernández, and N. Estrada-Saldívar. 2022. Stony coral tissue loss disease decimated Caribbean coral populations and reshaped reef functionality. Communications Biology 5:1–10.

Anderson, D. R. 2007. Model based inference in the life sciences: a primer on evidence. Springer Science & Business Media.

Benjamini, Y., and Y. Hochberg. 1995. Controlling the false discovery rate: a practical and powerful approach to multiple testing. Journal of the Royal statistical society: series B (Methodological) 57:289–300.

Brandt, M. E., R. S. Ennis, S. S. Meiling, J. Townsend, K. Cobleigh, A. Glahn, J. Quetel, V. Brandtneris, L. M. Henderson, and T. B. Smith. 2021. The Emergence and Initial Impact of Stony Coral Tissue Loss Disease (SCTLD) in the United States Virgin Islands. Frontiers in Marine Science:1105.

Clark, A. S., S. D. Williams, K. Maxwell, S. M. Rosales, L. K. Huebner, J. H. Landsberg, J. H. Hunt, and E. M. Muller. 2021. Characterization of the Microbiome of Corals with Stony Coral Tissue Loss Disease along Florida’s Coral Reef. Microorganisms 9:2181.

Dahlgren, C., V. Pizarro, K. Sherman, W. Greene, and J. Oliver. 2021. Spatial and Temporal Patterns of Stony Coral Tissue Loss Disease Outbreaks in The Bahamas. Frontiers in Marine Science:767.

Dobbelaere, T., D. M. Holstein, E. Muller, L. G. Gramer, L. McEachron, S. D. Williams, and E. Hanert. 2022. Connecting the dots: Transmission of stony coral tissue loss disease from the Marquesas to the Dry Tortugas. Frontiers in Marine Science 9.

Enochs, I., and G. Kolodziej. 2018. Ultraviolet deactivation of coral disease lesions. Florida DEP. Miami, FL.

Estrada-Saldívar, N., A. Molina-Hernández, E. Pérez-Cervantes, F. Medellin-Maldonado, F.J. González-Barrios, and L. Alvarez-Filip. 2020. Reef-scale impacts of the stony coral tissue loss disease outbreak. Coral Reefs 39:861–866.

Estrada-Saldívar, N., B.A. Quiroga-García, E. Pérez-Cervantes, O. O. Rivera-Garibay, and L. Alvarez-Filip. 2021. Effects of the stony coral tissue loss disease outbreak on coral communities and the benthic composition of Cozumel reefs. Frontiers in Marine Science 8:632777.

Evans, J. S., V. J. Paul, and C. A. Kellogg. 2022. Biofilms as potential reservoirs of stony coral tissue loss disease. Frontiers in Marine Science 9.

Hawthorn, A. C., M. Dennis, Y. Kiryu, J. Landsberg, E. Peters, and T. M. Work. 2024. Stony coral tissue loss disease (SCTLD) case definition for wildlife. U.S. Geological Survey Techniques and Methods, book 19, chapter 1.

Hayes, N. K., C. J. Walton, and D. S. Gilliam. 2022. Tissue loss disease outbreak significantly alters the Southeast Florida stony coral assemblage. Frontiers in Marine Science 9.

Heres, M. M., B. H. Farmer, F. Elmer, and H. Hertler. 2021. Ecological consequences of stony coral tissue loss disease in the Turks and Caicos Islands. Coral Reefs 40:609–624.

Hothorn, T., F. Bretz, and P. Westfall. 2008. Simultaneous inference in general parametric models. Biometrical Journal: Journal of Mathematical Methods in Biosciences 50:346–363.

Kramer, P., R. L, and L. J. 2025. Map of stony coral tissue loss disease outbreak in the Caribbean. https://www.agrra.org ArcGIS online.

Landsberg, J. H., Y. Kiryu, E. C. Peters, P. W. Wilson, N. Perry, Y. Waters, K. E. Maxwell, L. K. Huebner, and T. M. Work. 2020. Stony coral tissue loss disease in Florida is associated with disruption of host–zooxanthellae physiology. Frontiers in Marine Science 7:576013.

Lenth, R. 2018. Package ‘lsmeans’. The American Statistician 34:216–221.

Meiling, S., E. M. Muller, T. B. Smith, and M. E. Brandt. 2020. 3D photogrammetry reveals dynamics of stony coral tissue loss disease (SCTLD) lesion progression across a thermal stress event. Frontiers in Marine Science 7:597643.

Meiling, S. S., E. M. Muller, D. Lasseigne, A. Rossin, A. J. Veglia, N. MacKnight, B. Dimos, N. Huntley, A. Correa, and T. B. Smith. 2021a. Variable species responses to experimental stony coral tissue loss disease (SCTLD) exposure. Frontiers in Marine Science 8:464.

Meiling, S. S., E. M. Muller, D. Lasseigne, A. Rossin, A. J. Veglia, N. MacKnight, B. Dimos, N. Huntley, A. M. Correa, and T. B. Smith. 2021b. Variable species responses to experimental stony coral tissue loss disease (SCTLD) exposure. Frontiers in Marine Science 8:670829.

Muller, E. M., C. Sartor, N. I. Alcaraz, and R. van Woesik. 2020. Spatial Epidemiology of the Stony-Coral-Tissue-Loss Disease in Florida. Frontiers in Marine Science 7:163.

Neely, K. 2018. Ex-Situ disease treatment trials. Florida DEP. Miami, FL.

Neely, K. L. 2020. Novel treatment options for SCTLD final report. Florida DEP.

Miami FL. Neely K.L., C. L. Lewis, K. S. Lunz, and L. Kabay. 2021a. Rapid Population Decline of the Pillar Coral Dendrogyra cylindrus Along the Florida Reef Tract. Frontiers in Marine Science 8.

Neely, K. L., K. A. Macaulay, E. K. Hower, and M. A. Dobler. 2020. Effectiveness of topical antibiotics in treating corals affected by Stony Coral Tissue Loss Disease. Peerj 8:e9289.

Neely, K. L., R. J. Nowicki, M. A. Dobler, A. A. Chaparro, S. M. Miller, and K. A. Toth. 2024. Too hot to handle? The impact of the 2023 marine heatwave on Florida Keys coral. Frontiers in Marine Science 11:1489273.

Neely, K. L., C. P. Shea, K. A. Macaulay, E. K. Hower, and M. A. Dobler. 2021b. Short- and Long-Term Effectiveness of Coral Disease Treatments. Frontiers in Marine Science 8.

Precht, W. F., B. E. Gintert, M. L. Robbart, R. Fura, and R. van Woesik. 2016. Unprecedented Disease-Related Coral Mortality in Southeastern Florida. Scientific Reports 6:31374.

R Core Team. 2022. R: A language and environment for statistical computing. R Foundation for Statistical computing, Vienna, Austria.

Rosales, S. M., A. S. Clark, L. K. Huebner, R. R. Ruzicka, and E. Muller. 2020. Rhodobacterales and Rhizobiales are associated with Stony Coral Tissue Loss Disease and its suspected sources of transmission. Frontiers in Microbiology 11:681.

Rosales, S. M., L. K. Huebner, J. S. Evans, A. Apprill, A. C. Baker, C. C. Becker, A. J. Bellantuono, M. E. Brandt, A. S. Clark, and J. Del Campo. 2023. A meta-analysis of the stony coral tissue loss disease microbiome finds key bacteria in unaffected and lesion tissue in diseased colonies. ISME communications 3:19.

Shilling, E. N., I. R. Combs, and J. D. Voss. 2021. Assessing the effectiveness of two intervention methods for stony coral tissue loss disease on Montastraea cavernosa. Scientific Reports 11:1–11.

Sing, T., O. Sander, N. Beerenwinkel, and T. Lengauer. 2005. ROCR: visualizing classifier performance in R. Bioinformatics 21:3940–3941.

Studivan, M. S., M. Baptist, V. Molina, S. Riley, M. First, N. Soderberg, E. Rubin, A. Rossin, D. M. Holstein, and I. C. Enochs. 2022. Transmission of stony coral tissue loss disease (SCTLD) in simulated ballast water confirms the potential for ship-born spread. Scientific Reports 12:19248.

Therneau, T. 2022. A package for survival analysis in R. R package version 3.4.

Therneau, T. 2024. A package for survival analysis in R. R package version 3.8-3.

Therneau, T. M., and P. M. Grambsch. 2000. Modeling Survival Data: Extending the Cox Model. Springer.

Veglia, A., K. Beavers, E. Van Buren, S. Meiling, E. Muller, T. Smith, D. Holstein, A. Apprill, M. Brandt, and L. Mydlarz. 2022. Alphaflexivirus genomes in stony coral tissue loss disease-affected, disease-exposed, and disease-unexposed coral colonies in the US Virgin Islands. Microbiology Resource Announcements 11:e01199–01121.

Walker, B. K., H. Noren, S. Buckley, and K. Pitts. 2021a. Optimizing stony coral tissue loss disease (SCTLD) intervention treatments on Montastraea cavernosa in an endemic zone. Frontiers in Marine Science 8:746.

Walker, B. K., N. Turner, A. Brunelle, H. Noren, and S. F. Buckley. 2021b. 2020-2021 coral ECA reef-building-coral disease intervention and prepration for restoration final report. Florida DEP. Miami, FL.

Williams, S., and E. Muller. 2025. Testing coral disease treatments across multiple species at several highly-infected coral reef parks. Final Report. National Park Service PMIS Agreement P19AC01005.

Work, T. M., T. M. Weatherby, J. H. Landsberg, Y. Kiryu, S. M. Cook, and E. C. Peters. 2021. Viral-Like Particles Are Associated With Endosymbiont Pathology in Florida Corals Affected by Stony Coral Tissue Loss Disease. Frontiers in Marine Science 8.

